# Sympathetic Nervous System Overactivation Induces Colonic Eosinophil-Associated Microinflammation and Contributes to the Pathogenesis of Irritable Bowel Syndrome

**DOI:** 10.1101/2024.07.22.604223

**Authors:** Shaoqi Duan, Hirosato Kanda, Feng Zhu, Masamichi Okubo, Taro Koike, Yoshiya Ohno, Toshiyuki Tanaka, Yukiko Harima, Kazunari Miyamichi, Hirokazu Fukui, Shinichiro Shinzaki, Yilong Cui, Koichi Noguchi, Yi Dai

## Abstract

**Objective:** Mucosal microinflammation is a characteristic clinical manifestation of irritable bowel syndrome (IBS), and its symptoms are often triggered by psychological stress. In the present study, we aimed to investigate the impact of early life stress-associated dysfunction of the sympathetic nervous system (SNS) on mucosal immune changes in the gastrointestinal tract (GI) and its contribution to IBS pathogenesis.

**Design:** We utilised a traditional animal model of IBS with maternal separation (MS) and evaluated colorectal hypersensitivity, immune alterations, and SNS activity in adult rats with MS. We conducted a series of experiments to manipulate peripheral SNS activity pharmacologically and chemogenetically to explore the interaction between SNS activity and GI events.

**Results:** The MS-induced IBS model exhibited visceral hypersensitivity and eosinophilic infiltration in the colonic mucosa, along with SNS overactivation. Degeneration of the SNS using 6-OHDA neurotoxin decreased eosinophil infiltration and visceral hypersensitivity in the MS model. Notably, specific chemogenetic activation of the peripheral SNS induced eosinophil infiltration in the intestinal mucosa through the noradrenergic signalling-mediated release of eotaxin-1 from mesenchymal cells.

**Conclusion:** This study highlights the critical role of SNS overactivation in eotaxin-1-driven eosinophil infiltration in the colon, leading to the development of visceral hypersensitivity in IBS. The results provide important insights into the mechanistic links among increased sympathetic activity, mucosal microinflammation, and visceral hypersensitivity in individuals with IBS, suggesting potential therapeutic approaches.

**What is already known on this topic:**

- A subgroup of patients with irritable bowel syndrome (IBS) presents with microinflammation in the gastrointestinal tract (GI).
- Early life stress is recognised as a major risk factor for the development of IBS in adulthood.
- Overactivation of the sympathetic nervous system (SNS) is frequently associated with IBS.

**What this study adds:**

- Maternal separation (MS) stress induces eosinophil-associated microinflammation in the colonic mucosa of adult rats.
- Inhibition of SNS activity suppresses eosinophil infiltration and mitigates visceral hypersensitivity in the MS model.
- Noradrenergic signalling within the peripheral sympathetic activation stimulates mesenchymal cells to release eotaxin-1, leading to substantial eosinophil-predominant immune alterations in the colon.

**How this study might affect research, practice, or policy:**

- Treatment with fibroblast-derived eotaxin-1 and targeting eosinophil-associated microinflammation could be a potential strategy to alleviate visceral pain in patients with IBS.
- The chemogenomic method specifically manipulates peripheral SNS and provides a valuable tool for future research.

## INTRODUCTION

Irritable bowel syndrome (IBS) is a prevalent functional gastrointestinal disorder (FGID) characterised by intermittent or persistent abdominal pain, diarrhoea, and constipation without apparent organ abnormalities ^1^. These recurring symptoms often impact the quality of life and are closely associated with mental stress. Although IBS affects approximately 15% of the global population ^2^, effective clinical treatment remains challenging owing to a limited understanding of its underlying pathophysiological mechanism.

The pathogenesis of IBS involves a complex interplay of genetic, environmental, and immunological factors ^3^. Various pathological mechanisms, including visceral hypersensitivity, altered intestinal motility, immune dysfunction, and dysbiosis, are implicated in IBS ^4^. Notably, some patients with IBS exhibit microinflammation (low-grade inflammation) in the gastrointestinal tract (GI) mucosa ^5^, challenging the conventional view that FGIDs lack organic abnormalities. Clinical evidence indicates that a considerable subgroup of individuals with IBS experiences microinflammation in the colonic mucosa, characterised by infiltration and activation of immune cells within the GI mucosa ^4–6^. Mast cells have garnered attention owing to their contribution to visceral hypersensitivity ^6–8^. Additionally, higher numbers of eosinophils, lymphocytes, and other plasma cells have been reported in the intestinal mucosae of patients with IBS than in those of healthy individuals ^5, 9, 10^. These reported variations in the types of immune cells contributing to microinflammation may be attributed to geographic differences in diet, genetic backgrounds, patient phenotypes, or biopsy sites ^11^. Despite numerous reports of increased immune cell numbers in the mucosa of patients with IBS, underlying mechanisms inducing alterations in the intestinal immune system remain unknown.

Psychosocial factors (particularly early life adverse events and traumatic stress) are strongly implicated in the pathophysiology of IBS. Specifically, individuals exposed to stress in early life are more likely to develop IBS during adulthood ^12, 13^. In a subset of patients with IBS who have experienced early life stress, the sympathetic nervous system (SNS) becomes more sensitive to stress, leading to sympathetic overactivation ^14, 15^. The mechanisms underlying the influence of psychosocial events on IBS have been investigated, with particular focus on the role of the hypothalamic-pituitary-adrenal (HPA) axis dysregulation, a neuroendocrine pathway ^16–18^. However, the contribution of the peripheral SNS remains unknown. Emerging evidence indicates an interaction between the SNS and the immune system ^19^. The SNS uses norepinephrine (NE) as its primary neurotransmitter to regulate communication between sympathetic neurons and immune cells. Generally, SNS activation suppresses immune responses ^20,21^. However, in certain cases, NE signalling induces immune cell accumulation in local tissues, subsequently enhancing immune responses ^22, 23^. Therefore, the SNS can exert both activating and inhibitory effects on immunity ^19^. The occurrence of both immune cell alterations and sympathetic perturbation in patients with IBS suggests a potential causal relationship between an overactive SNS and microinflammation, contributing to IBS pathogenesis.

We have previously reported that maternal separation (MS) stress induces eosinophil-associated microinflammation in the gastroduodenal mucosa, which is linked with increased gastric hypersensitivity to distention ^24^. Mucosal eosinophils are key neuroimmune players in regulating GI function and are implicated in brain-gut interaction disorders ^25^. Herein, we used the MS model ^26^ to mimic the eosinophil-associated microinflammation observed in patients with IBS and examined the influence of SNS overactivity on immune alterations in the GI. Specifically, we introduced two chemogenomic approaches to manipulate SNS activity, which enabled the manipulation of peripheral sympathetic activity without the confounding effects of glucocorticoids.

## MATERIALS AND METHODS

### Animals and study approval

B6.Cg-Tg (Dopamine beta-hydroxylase [DBH]-cre) 9-9Koba/KobaRbrc (RBRC01492) mice were acquired from Professor Kobayashi ^27^, RIKEN BRC, through the National BioResource Project of MEXT/AMED, Japan. Cocaine- and amphetamine-regulated transcript protein (Cartpt)-Cre mice (accession No. CDB0267E) lines (listed in https://large.riken.jp/distribution/mutant-list.html) were generated via CRISPR/Cas9-mediated knock in zygotes as reported previously ^28^. Fourteen-day pregnant and adult (7 weeks old) Sprague-Dawley rats were purchased from Japan SLC Inc. and housed in a standard laboratory with free access to water and food. Male litters were used in the study. Experimental protocols were designed to minimise the number of animals used and were approved by the Committee on Animal Research at Hyogo Medical University (#2021-22, 2021-18, and 2021-04).

### Human endoscopic biopsy specimens

Patients diagnosed with IBS were identified based on the Roma IV criteria ^1^. None of the participants had taken any anti-inflammatory medications. Colorectal biopsy specimens were obtained from healthy controls (n = 3 for female, n = 1 for male) and patients (n = 2 for female, n = 2 for male) during endoscopy and used for immunohistochemical staining. All patients with IBS and controls were recruited from Hyogo Medical University Hospital, Japan. This study was performed in accordance with the Declaration of Helsinki and was approved by The Ethics Review Board of Hyogo Medical University (No. 4706). All participants provided written informed consent. Patients were not involved in the design, or conduct, or reporting, or dissemination plans of our research.

### MS model

The MS rat model was established as described previously ^24^. Briefly, newborn pups were randomly divided into the control and MS groups. MS pups were individually placed in disposable paper cups (5 cm in diameter) for 3 h per day for 14 days. After separation, the pups were returned to their mother’s cage; control pups remained with their mothers continuously. The rats were reared for up to 7 weeks for subsequent experiments.

### Adeno-associated virus (AAV) infection

AAV PHP.S-hSyn-DIO-hM3D-mCherry (1 × 10^13^ viral genomes/mL) was injected into the retro-orbital sinuses of 4-week-old DBH-cre mice. AAV8 hSyn-DIO-hM3D (Gq)-mCherry (2.1 × 10^13^ viral genomes/mL; Addgene #44361) or AAV8 hSyn-DIO-mCherry (2.3 × 10^13^ viral genomes/mL; Addgene #50459) was injected into the 8–12th thoracic spinal cord of 4-week-old Cartpt-Cre mice. The detailed experimental procedure is provided in Supplemental Methods.

### Colon lamina propria cell harvest and flow cytometry

Colon tissues were collected 6 h after NE colorectal infusion from wild-type mice or after 5 days of continuous clozapine N-oxide (CNO) stimulation from DBH-cre mice. Colonic samples were digested into a single-cell suspension and incubated with appropriate antibodies for flow cytometry. Adhesion signals were initially excluded during flow cytometry analysis, and approximately 50,000 events were gated to live cells using 4’,6-diamidino-2-phenylindole (DAPI) for each sample. Subsequently, the gates were set on anti-CD45 positive events, followed by gating for specific immune cells. FlowJo software (BD Biosciences, Franklin Lakes, NJ, USA) Version 10.8.1 was used for data analysis. Detailed processing steps and antibodies used are provided in Supplemental Methods.

### Immunohistochemistry (IHC)

IHC was performed as described previously ^24^. Brain, superior cervical ganglion (SCG), dorsal root ganglion (DRG), and GI tissues (3–5 cm from the anus of rats and whole colon of mice) were dissected and fixed with paraformaldehyde or acetone. Subsequently, tissue sections were prepared for antibody incubation. The primary and secondary antibodies used are listed in Supplemental Tables 1 and 2, and further experimental details are described in Supplemental Methods.

### Enzyme-linked immunosorbent assay (ELISA)

Eotaxin-1 release from CCD-18co cell lines following NE stimulation (0, 1, and 10 μM NE after 1, 3, or 6 h of incubation) was measured using the human CCL11/Eotaxin kit (R&D Systems, Minneapolis, MN, USA) according to the manufacturer’s protocol. Blood NE concentrations of MS model rats and DBH-cre mice were detected using an NE ELISA kit (LifeSpan Biosciences, Lynwood, WA, USA) following the manufacturer’s instructions. Details are presented in Supplemental Methods.

### Reverse transcription PCR (RT-PCR)

Samples of CCD-18co cells after NE stimulation (0, 1, and 10 μM NE after 1, 3, or 6 h of incubation), CD34^+^PDGFRα^+^ cells sorted via FACS, and rat colonic tissues were collected for RT-PCR as reported previously ^24^. Sample collection is detailed in Supplemental Methods. The primers used are listed in Supplemental Table 3.

### Colorectal infusion

Wild-type rats and mice were anaesthetised with 2% isoflurane for NE stimulation and eotaxin-1 colorectal infusion experiments. A plastic oral gavage needle was slowly inserted 3.5 cm into the colon through the anus, and NE (0.5 mg/mL, 1 mL/kg) or eotaxin-1 (1 mg/mL, 1 ml/kg) was then infused into the colon.

### Pharmacologic agents

6-OHDA, CNO, SB328437, phentolamine, and propranolol were administered intraperitoneally. The dose and time course are detailed in Supplemental Methods.

### Data analysis

Statistical analyses were performed using Microsoft Excel and GraphPad Prism 9.0 (GraphPad Software, Inc., La Jolla, CA, USA). Statistical methods used in the study are indicated in figure legends. All data are reported as mean ± standard error. P-values < 0.05 were considered statistically significant (* P < 0.05, ** P < 0.01, and *** P < 0.001).

## RESULTS

### MS stress-induced IBS model exhibits eosinophil infiltration and SNS overactivation

We used a traditional MS stress-induced IBS model to induce colonic microinflammation and visceral hypersensitivity. Adult MS rats exhibited visceral hypersensitivity, as evidenced by increased visceromotor response (VMR) at 40 and 60 mmHg, compared with control rats (Figure 1A, B). The threshold of the colorectal distention (CRD) response did not differ significantly between MS and control groups (Figure 1C), indicating that MS stress induced visceral hyperalgesia rather than allodynia in adulthood. These data confirm that MS stress induces visceral hypersensitivity to CRD in adulthood, a characteristic symptom of IBS.

**Figure 1.**
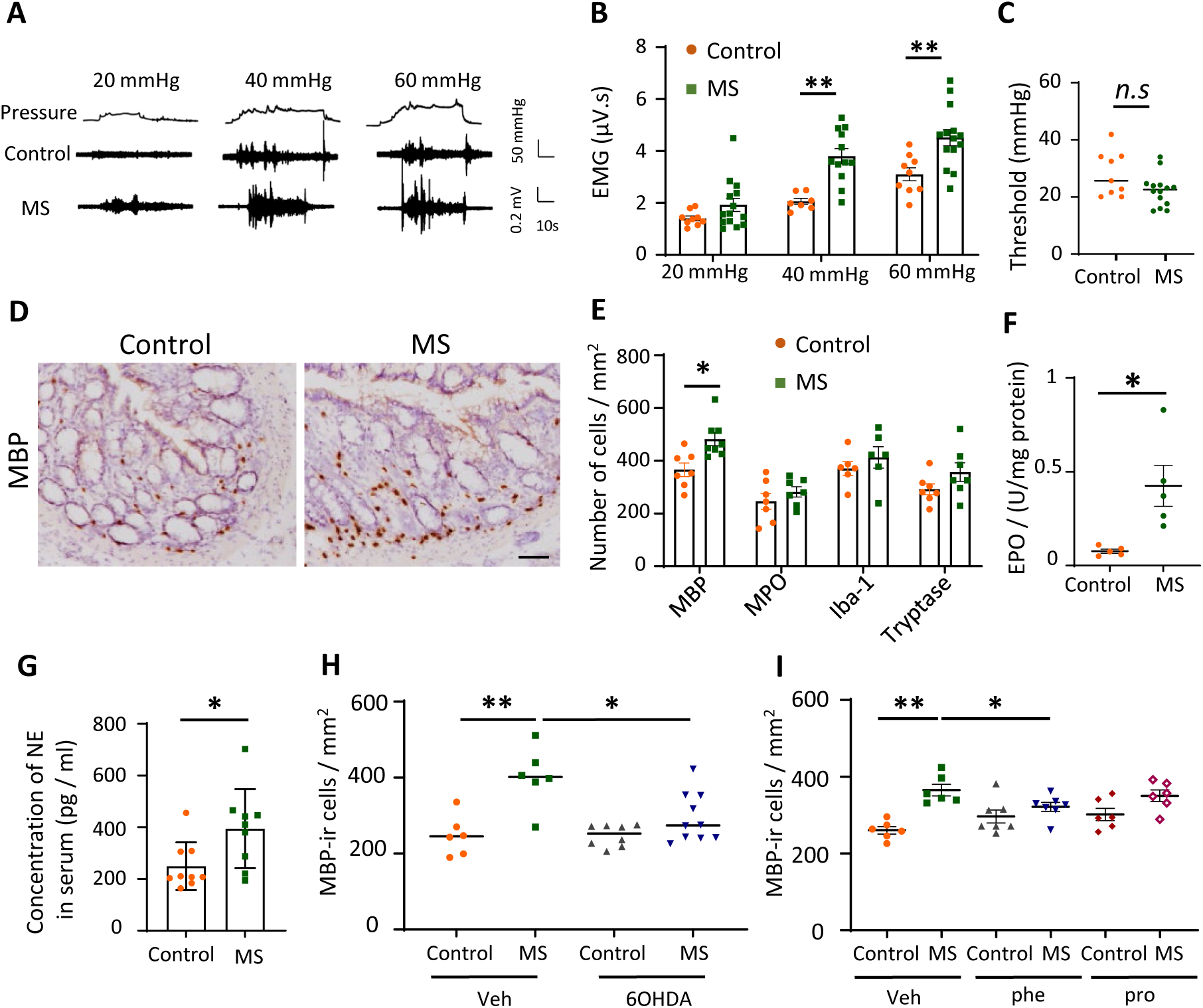
Maternal separation (MS) stress-induced irritable bowel syndrome (IBS) model exhibits eosinophil infiltration and overactivation of the sympathetic nervous system (SNS). (A) Colorectal distention pressures of 20, 40, and 60 mmHg were applied to rats in both the control and the MS groups (top). Electromyography (EMG) traces of visceral motor response (VMR) to colorectal distention (CRD) at different pressures are shown for the control (middle) and MS (bottom) groups. (B, C) Bar graphs and dot plots represent the data of VMR to colorectal distention at different pressures (B) and the threshold (C), respectively (n = 10–14 rats in each group). (D) Representative images of colonic tissue stained for major basic protein-immunoreactive (MBP-ir) cells in the colon of the control and MS groups. Brown dots indicate MBP-ir cells. Scale bar, 50 μm. (E) Bar graphs represent the number of immune cells in the colonic mucosa of the control and MS groups, including MBP, myeloperoxidase (MPO), and ionised calcium-binding adapter molecule 1 (Iba-1) ir cells (n = 6-7 in each group). (F) Eosinophil peroxidase (EPO) activity in the colon of the control and the MS groups (n = 5 rats in each group). (G) Bar graphs show the norepinephrine (NE) concentration in the serum of the control and MS groups (n = 9 rats in each group). (H) Scatter dot plot data showing the number of MBP-ir cells in the control and MS groups in the presence or absence of 6-OHDA (n = 6–10 rats in each group). (I) Scatter dot plot showing the number of MBP-ir cells in the control and MS groups treated or untreated with adrenergic receptor (AR) antagonists (Veh: vehicle; Phe: phentolamine; Pro: propranolol) (n = 6 rats in each group). Data are represented as mean ± SEM. Two-way ANOVA with Bonferroni post hoc test was used for (B, H, I), and an unpaired *t*-test was used for (C, E, F, G). *P < 0.05, **P < 0.01. n.s; P > 0.05.

Immune alteration has been reported in patients with FGIDs and is associated with clinical symptom severity ^4^. The MS group exhibited a significantly higher number of major basic protein (MBP)-immunoreactive (-ir) cells in the colonic tissue than the control group (Figure 1D, E). There was a trend towards a significant increase in the number of tryptase-ir cells (p = 0.138). The number of myeloperoxidase (MPO)-ir and ionised calcium-binding adapter molecule 1(Iba-1)-ir cells did not differ significantly between the control and MS groups (Figure 1E, Supplemental Figure S1A). Additionally, the MS group exhibited no obvious morphological differences in the colonic mucosal tissues (Supplemental Figure S1B). The relative mRNA expression of MBP (*Prg3*) rather than that of tryptase (*Tpsab1*) was increased in the colon of the MS model (Supplemental Figure S1C). These findings indicate that eosinophils, rather than neutrophils, macrophages, or mast cells, predominantly infiltrate the lamina propria of the colon in MS rats. Additionally, increased eosinophil peroxidase activity was observed in the colon of the MS group, indicating an increased activation status of eosinophils (Figure 1F).

Neuroimmune interactions are important in the immune response ^19^. Given the common feature of increased SNS activity in IBS ^13, 29^, we quantified SNS activity in the MS group by monitoring the NE serum concentration, which was significantly higher than that in the control group (Figure 1G). These data indicated that MS rats developed eosinophil infiltration and exhibited an overactive SNS. To investigate whether the overactive sympathetic signal contributes to eosinophil infiltration in the MS model, we blocked sympathetic activity using 6-OHDA (a neurotoxic compound selective for dopaminergic and noradrenergic neurones). Intraperitoneal injection of 6-OHDA (7 days after treatment) significantly suppressed eosinophil infiltration in the MS group (Figure 1H). Additionally, treatment with phentolamine, an alpha-adrenergic receptor (AR) antagonist, for seven continuous days significantly suppressed eosinophil infiltration compared with that in the vehicle-treated MS group. Moreover, treatment with phentolamine and propranolol did not show significant eosinophil infiltration in the MS group (Figure 1I). These data indicate that sympathetic signalling plays a role in mucosal eosinophil migration in MS rats.

### Genetic activation of peripheral SNS induces eosinophil infiltration in the colon

To further investigate the regulatory role of sympathetic activity in eosinophil migration, we employed a genetic manipulation technique to control peripheral sympathetic signals without affecting adrenal corticoid function. Using AAV PHP.S-hSyn-DIO-hM3D-mCherry^30^, we modulated peripheral SNS activity in DBH-cre mice (Figure 2A). We quantified the infection rate in the SCG, where AAV infected 24.8 ± 1.14% of neurones in a cre-dependent manner, without any detectable infection in the central nervous system (CNS) and somatosensory system (Supplemental Figure S2A-C). This strategy enabled the targeted activation of DBH-cre positive cells to modulate peripheral SNS activity, evidenced by higher serum NE concentrations after continuous CNO stimulation for 5 days (Figure 2B). Moreover, treatment with CNO reduced the excretion of faecal pellets from the colon, supporting enhanced SNS activity (Figure 2C). As AAV-PHP.S primarily infects the peripheral nervous system rather than the CNS, we could examine the effect of peripheral SNS activity on immune alterations without the influence of the HPA axis.

**Figure 2.**
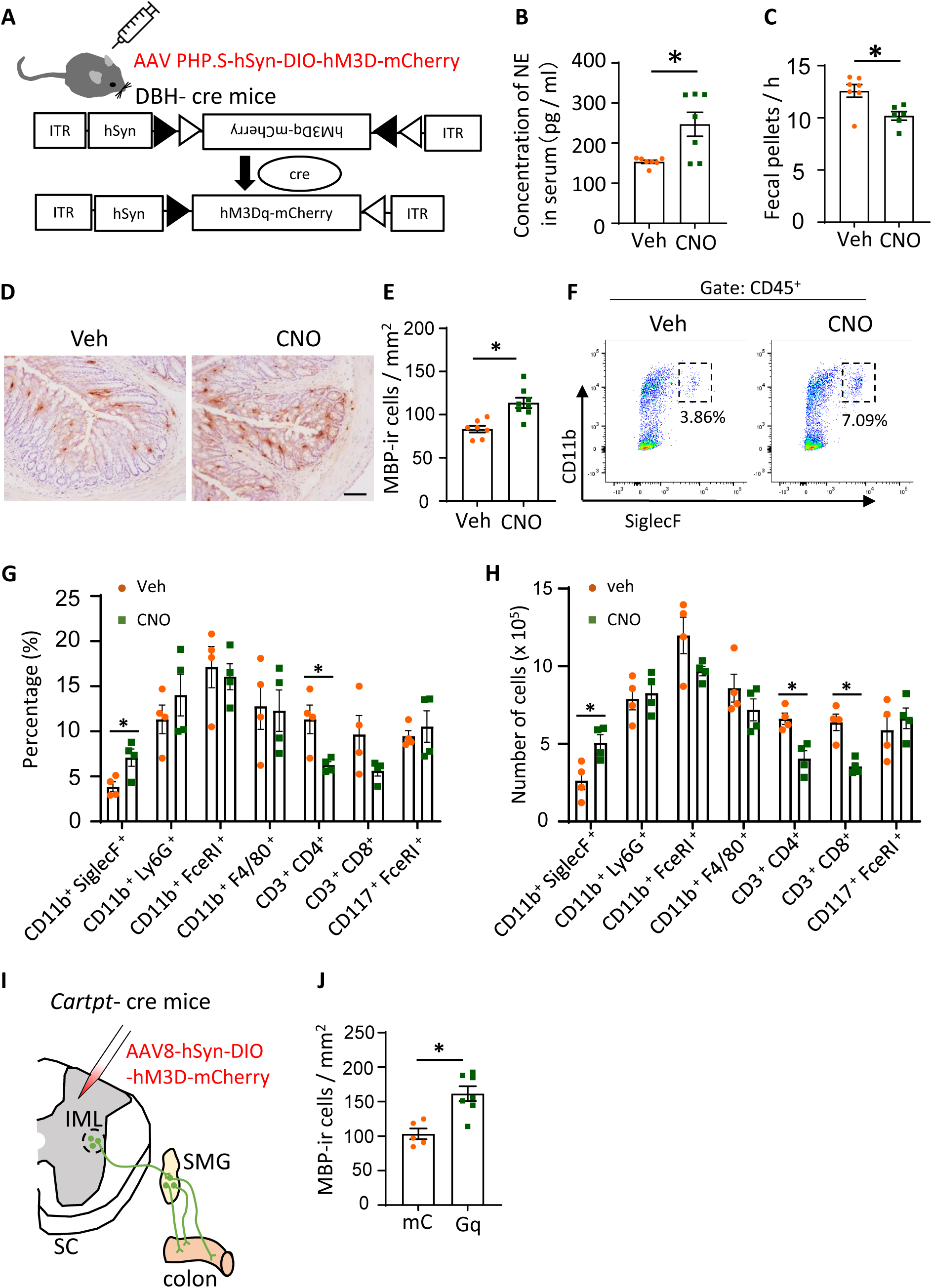
Genetic activation of peripheral DBH-cre cells induces eosinophil infiltration in the colon. (A) Schematic diagram illustrating the AAV-PHP.s-hSyn-DIO-hM3D construct expressing hM3D in DBH-cre mice to selectively modify peripheral DBH-positive cells. (B-C) Clozapine-*N*-oxide (CNO) treatment increases norepinephrine (NE) concentration in the serum (B) and decreases faecal pellet output from the colon (C) (n=7 in each group). (D) Immunohistochemistry images showing major basic protein-immunoreactive (MBP-ir) cells in the colorectal tissues of vehicle (Veh) and CNO-treated DBH-cre mice. Scale bar, 50 μm. (E) Bar graphs showing the number of MBP-ir cells in the colonic tissues of Veh- and CNO-treated DBH-cre mice. (n = 7 in each group). (F) Flow cytometry gated CD45^+^CD11b^+^SiglecF^+^ cells as eosinophils in the Veh and CNO-treated mice. (G) Bar graph showing the percentage of immune cells and (H) absolute cell number of immune cells in the Veh and CNO groups (n = 4 in each group, percentage in CD45+ cells). (I) Schematic diagram showing the strategy of specifically modulating the sympathetic nervous system (SNS) innervating the colon. (J) Bar graphs showing MBP-ir cell counts in the mCherry (mC) and Gq groups of *Cartpt-cre* mice after CNO treatment. Data are presented as mean ± SEM, with statistical analysis performed using the nonparametric Mann-Whitney test. *P < 0.05.

To investigate immune alterations in the colon with chemogenetic modulation of peripheral sympathetic activity, we initially examined the distribution of eosinophils in colorectal tissue via immunostaining. The CNO group showed a significant increase in the MBP-ir cell count in the lamina propria when compared with the vehicle group (Figure 2D, E), exhibiting a pattern similar to that in the MS model. To confirm the immune alteration specifically, we conducted flow cytometry on a single-cell suspension prepared from the colon. We first gated CD45+ cells as immune cells (Supplemental Figure S2D) and then used specific cell surface markers to identify target immune cells. The ratio and absolute cell number of colonic eosinophils (CD11b^+^ SiglecF^+^) significantly increased in the CNO group. No significant differences were observed in neutrophils (CD11b^+^ Ly6G^+^), basophils (CD11b^+^ FcεRI^+^), macrophages (CD11b^+^ F4/80^+^), or mast cells (CD117^+^ FcεRI^+^) between the CNO and vehicle-treated groups. The ratio of CD4^+^ T cells decreased after CNO treatment (Figure 2F, G, and H). Treatment with CNO did not significantly alter the proportion and number of any immune cells in naïve mice (Supplemental Figure S2C, D). The percentage of basophils (CD11b^+^ FcεRI^+^) was higher than the basal value under normal conditions, according to the FACS data, which could be attributed to the limitation of the FcεRI^+^ antibody recognising other cell types when used in rodents ^31^.

Although chemogenetically activating peripheral SNS in DBH-cre mice could exclude the effect from the CNS, the peripheral sympathetic nerve-innervating organs (including the adrenal medulla) would be activated following CNO injection. Therefore, we used Cartpt-cre mice with the AAV8-hSyn-DIO-hM3D-mcherry infection to selectively control the targeted preganglionic spinal neuron-CG/SMG-gastrointestinal outflow (Figure 2I). This system enabled precise manipulation of sympathetic activity in the gut (for detail see^28^). Following CNO treatment, the eosinophil count significantly increased in the colon of the Gq group compared with that in the colon of the mCherry group (Figure 2J), indicating that activation of colon-innervating sympathetic nerves can efficiently recruit eosinophils to the colon.

### NE colorectal infusion induces eosinophil infiltration into the colon

To further investigate the mechanisms involved in eosinophil recruitment mediated by the SNS, we performed NE colorectal infusion and examined the distribution of immune cells. Six hours after NE colorectal infusion, lamina propria cells were collected from the colon of mice for flow cytometric analysis. Following NE infusion in the colonic tissue, we detected a specific increase in the percentage and absolute cell number of eosinophils (relative to CD45^+^ cells) (Figure 3A–C). Notably, there were no significant increases in other types of immune cells. Consistently, IHC staining revealed a substantial increase in the number of MBP-ir cells after NE treatment, whereas neutrophils, macrophages, and mast cells showed no changes in the colon (Supplemental Figure S3A, B).

**Figure 3.**
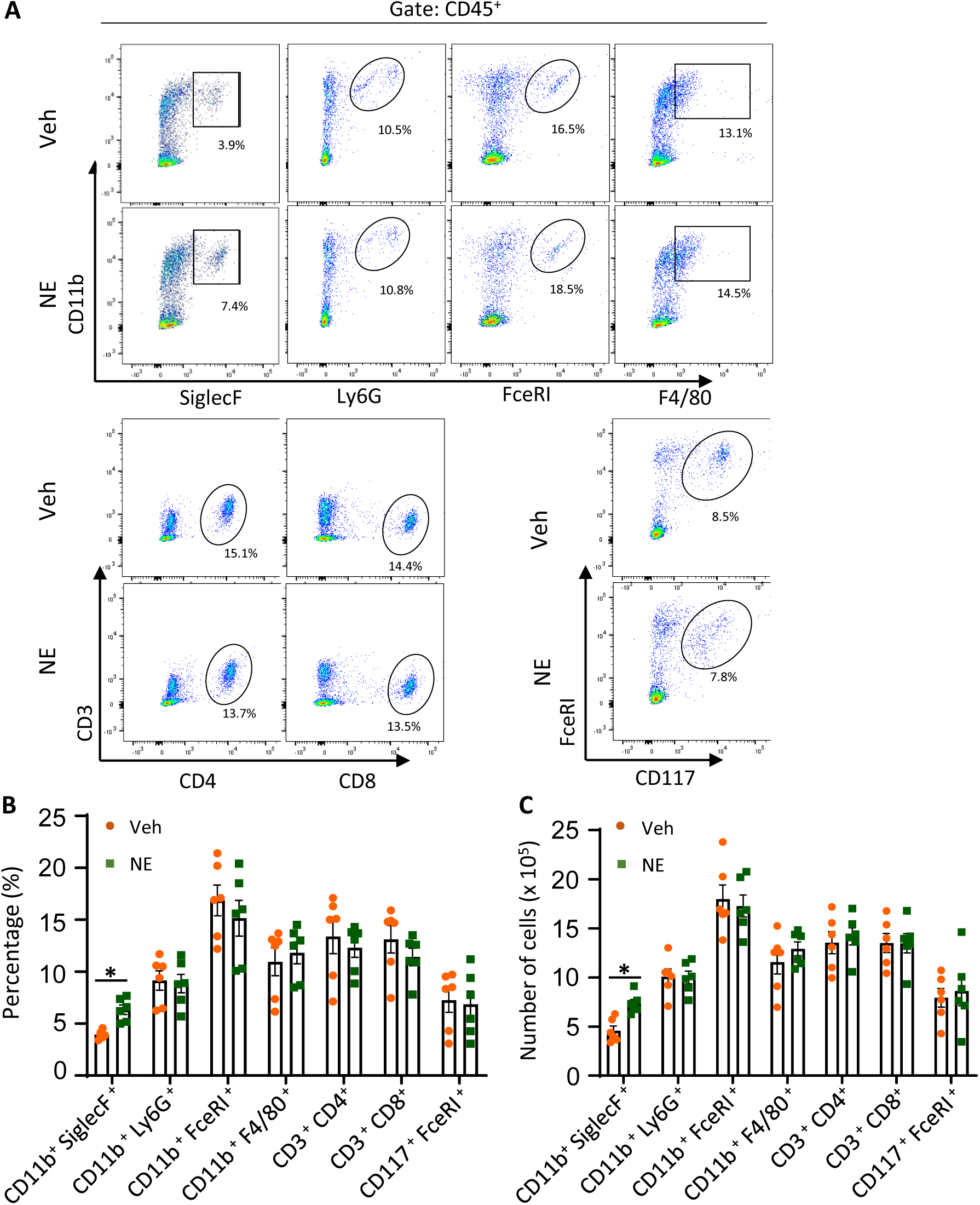
Norepinephrine (NE)-colorectal infusion induces eosinophil infiltration in the colon. (A) Flow cytometry analysis of immune cells gated in both vehicle (Veh) and NE-treated mice. (B, C) Bar graphs show the percentage (B) and absolute cell number (C) of immune cell alterations in the Veh- and NE-treated mice. (n= 6 in each group, the percentage in CD45+ cells). Data are represented as mean ± SEM, with statistical analysis performed using the nonparametric Mann-Whitney test. *P < 0.05.

To elucidate the role of ARs in eosinophil infiltration, we treated rats with two pan-AR antagonists (phentolamine and propranolol) for 1 h before NE infusion. Consistent with the findings observed in MS rats (Figure 1I), treatment with phentolamine significantly reduced eosinophil infiltration, whereas propranolol induced a trend towards a significant decrease in the number of eosinophils (p = 0.12). Meanwhile, there was no significant difference in the MBP-ir cell count between the vehicle- and propranolol-treated groups (Supplemental Figure S3C). These results, along with the data from MS rats (Figure 1I), suggest that both alpha-AR and beta-AR play a role in regulating eosinophil migration in the colonic mucosa.

### Eotaxin-1 mediates NE-induced eosinophil infiltration

As chemoattractants and cytokines play crucial roles in eosinophil recruitment to tissues ^32^, we identified specific cytokines and chemokines involved in NE-induced eosinophil infiltration. We examined the expression of various genes (*Ccl11*, *Ccl24*, *Ccl26, Il33,* and *Il5*) associated with eosinophil infiltration after NE infusion into the colon of rats. Eotaxin-1 (*Ccl11*) was significantly upregulated in the colon 6 h after NE infusion, whereas expression levels of *Ccl24* and *Il33* were unaltered, and the expression of *Ccl26* and *Il5* was undetectable in the colonic mucosa (Figure 4A, B). Correspondingly, the expression of eotaxin-1-ir cells significantly increased from 3 h after NE colorectal infusion (Figure 4C, D). In a parallel experiment, we detected no differences in the expression of *CCL11, IL5,* and *IL33* in the bone marrow (Supplemental Figure S3D) between the control and MS group, indicating that eosinophil accumulation is restricted to the colon rather than systemic circulation.

**Figure 4.**
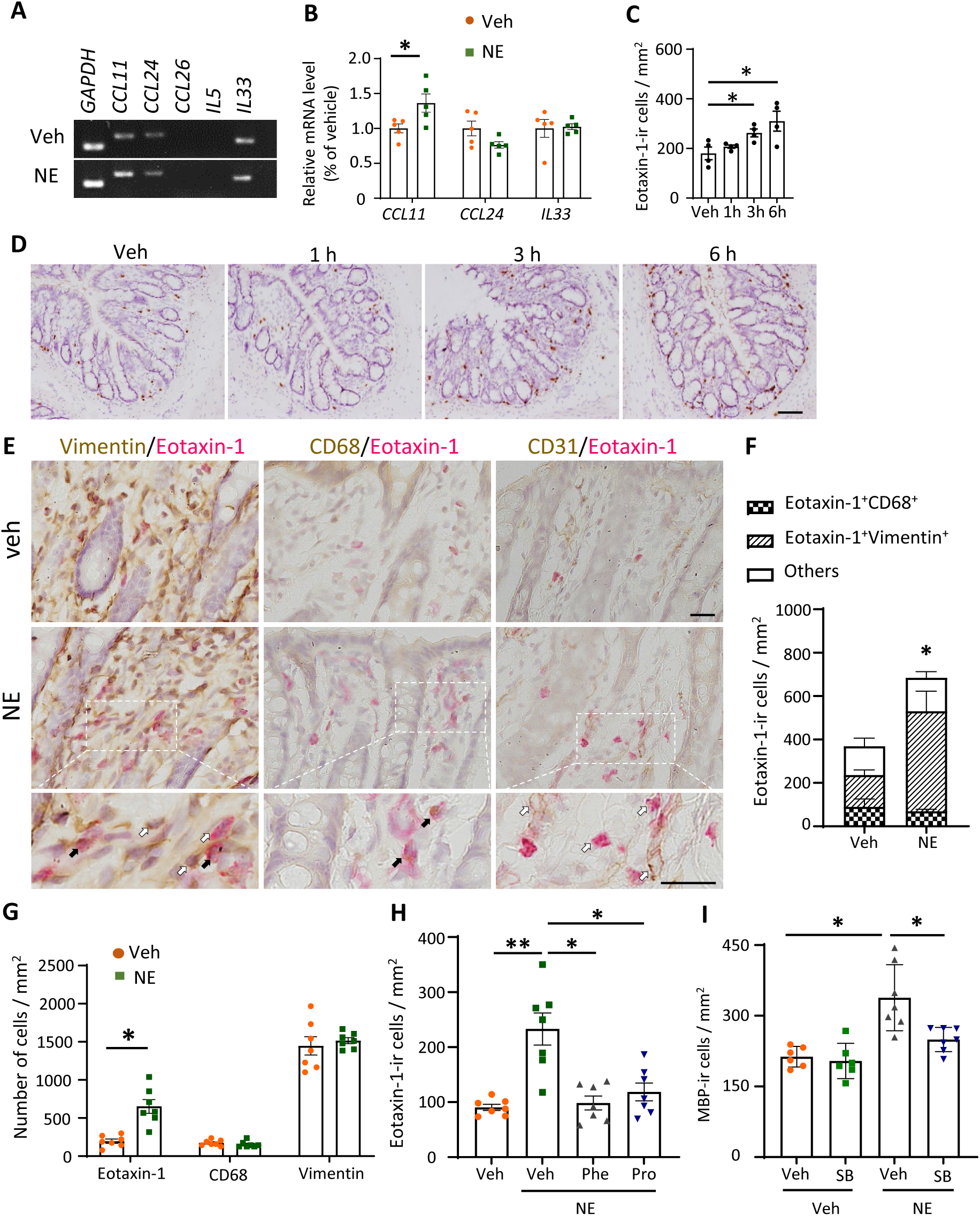
Eotaxin-1 contributes to the norepinephrine (NE)-induced eosinophil infiltration in the colon. (A) PCR analysis of the expression of eosinophil recruitment genes (*Ccl11, Cccl24, Ccl26, Il5,* and *Il33*) in the colons of vehicle (Veh)- and NE-treated rats 6 h after NE treatment. (B) Bar graphs showing relative changes in gene expression between the colons of Veh- and NE-treated rats (n = 5 in each group). (C) Summary data of the number of eotaxin-1-ir cells at different time points after NE treatment. *P < 0.05, vs. vehicle (Veh). (D) Immunohistochemistry images showing eotaxin-1 expression in the colon 6 h after NE treatment. Scale bar, 50 μm. (E) Immunohistochemistry images showing co-expression of eotaxin-1 with vimentin, CD68, and CD31 in the colon of Veh- and NE-treated rats, respectively. Enlarged images of the white dotted frame are shown at the bottom. Black arrows indicate double-labelled cells, and white arrows indicate single-labelled cells. Scale bar, 20 μm. (F) Bar graphs showing the eotaxin-1-ir cells in the Veh and NE stimulation groups (n =7 in each group). * P < 0.05, eotaxin-1-ir cells NE vs Veh. (G) Bar graphs showing data of eotaxin-1 cells, CD68 cells, and vimentin cells in the Veh- and NE-treated rats (n =7 in each group). (H) Bar graphs showing the number of eotaxin-1-ir cells with or without phentolamine (Phe) and propranolol (Pro) administration in Veh- and NE-treated rats (n= 7 in each group). (I) Bar graphs showing the number of MBP-ir cells with or without SB328437 administration in Veh- and NE-treated rats (n= 6 rats in each group). Data are presented as mean ± SEM. A one-way ANOVA was used for the analysis shown in (C, H), a two-way ANOVA with Bonferroni post hoc test was used for the analysis shown in (I), and an unpaired *t*-test was used for the analysis shown in (B, F, G). *P < 0.05, **P < 0.01.

Eotaxin-1 can be released by epithelial cells, endothelial cells, macrophages, and mesenchymal cells ^33, 34^. To identify the involvement of eotaxin-1-releasing cells after NE infusion, we performed double immunostaining of eotaxin-1 with CD68, vimentin, and CD31, representing markers of macrophages, mesenchymal cells, and endothelial cells, respectively (Figure 4E). In the vehicle-treated group, 46.2 ± 4.5% and 51.1 ± 3.2% of eotaxin-1-ir cells co-expressed CD68 and vimentin, respectively, while almost no eotaxin-1-ir cells co-expressed CD31. In the NE-treated group, the percentage of eotaxin-1-ir cells significantly increased. Double immunostaining indicated that 19.6 ± 3.7% and 74.5 ± 3.5% of eotaxin-1 cells co-expressed CD68 and vimentin, respectively (Figure 4F). Colonic NE infusion did not alter the total number of macrophage and mesenchymal cells (Figure 4G). These data suggest that NE colonic infusion upregulates eotaxin-1 predominantly in mesenchymal cells rather than other cell types.

To identify ARs involved in this regulation, we pre-treated rats with alpha- or beta-AR antagonists 1 h before NE stimulation. Treatment with either the alpha-AR antagonist, phentolamine or beta-AR antagonist propranolol inhibited the upregulation of eotaxin-1 (Figure 4H). Administration of the eotaxin-1 receptor inhibitor SB328437 significantly inhibited NE-induced eosinophil infiltration in the colonic mucosa (Figure 4I). These findings suggest that NE induces eosinophilic infiltration in the colon by upregulating eotaxin-1 in mesenchymal cells.

Furthermore, we ascertained whether chemogenetic activation of the SNS results in increased eotaxin-1 expression. The data from the selective activation of the SNS-gastrointestinal pathway in Cartpt-cre mice infected with AAV8-hSyn-DIO-hM3D-mcherry demonstrated a significant increase in eotaxin-1 expression in the colon. This suggests that SNS activity contributes to the increased eotaxin-1 expression (Supplemental Figure S4A, B).

### NE stimulates CCD-18co cells to release eotaxin-1

Intestinal fibroblasts, the main components of prototypical mesenchymal cells, share the same immunophenotypic properties ^35^. To confirm eotaxin-1 release from mesenchymal cells, we cultured CCD-18co cells, a non-malignant human fibroblast cell line originating from normal colon tissue (Figure 5A). Stimulation with 10 μM NE significantly upregulated the expression of *hCCL11* from 3 h onwards, whereas low concentrations and short treatment times did not affect expression (Figure 5B, C). Using ELISA, we found that NE stimulation resulted in a 13.8-fold increase in the protein level of eotaxin-1 in the culture supernatants at 3 h, which further increased to 35.7 times 6 h after NE stimulation (Figure 5D). These data demonstrate that human intestinal fibroblasts can release eotaxin-1 after NE stimulation.

**Figure 5.**
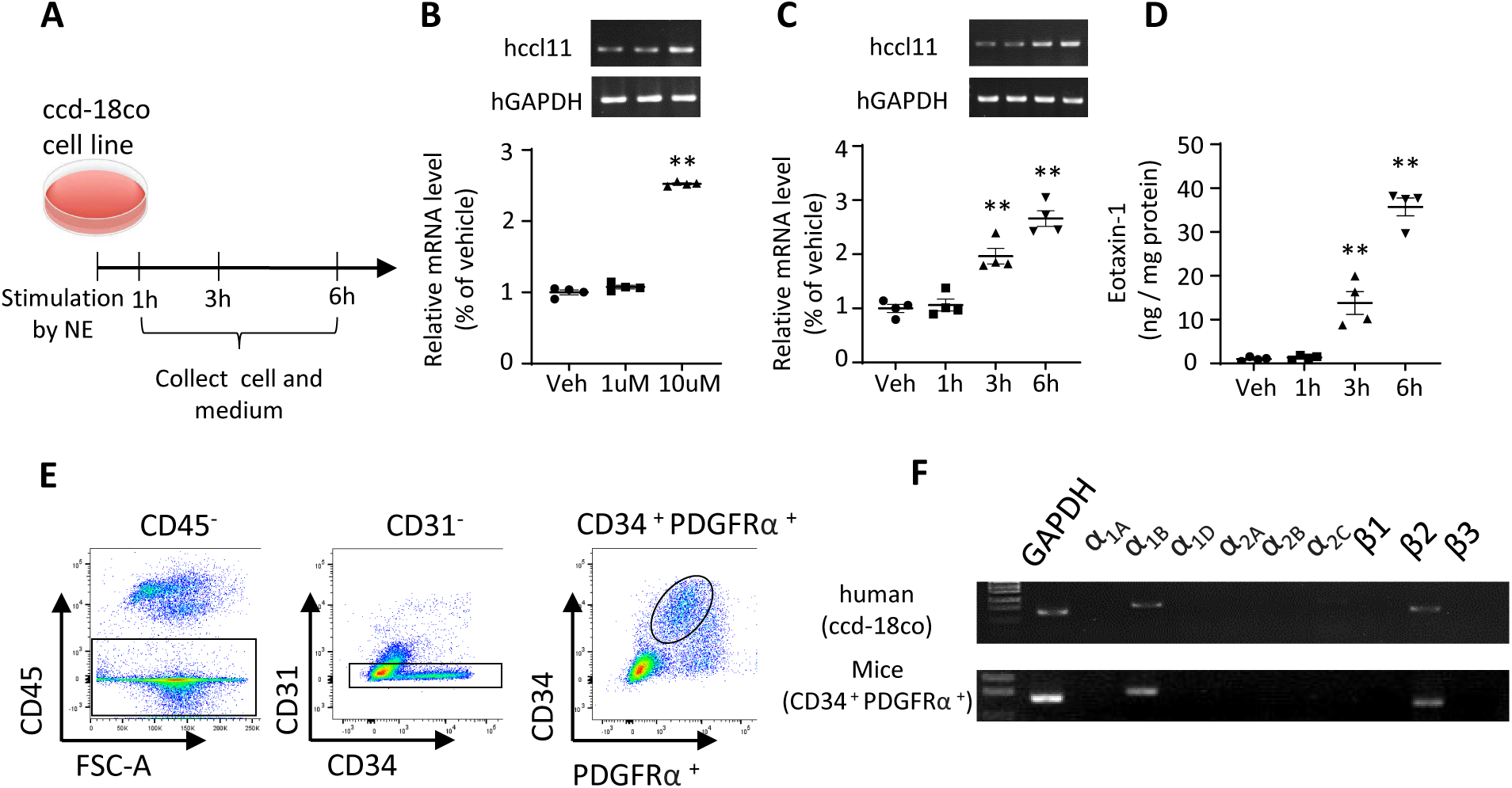
Eotaxin-1 is released from fibroblasts in response to norepinephrine (NE) stimulation. (A) The schematic diagram represents the protocols of NE treatment and sample collection. (B) Images and graphs show PCR bands and relative mRNA levels of *Hccl11* at different NE concentrations (vehicle [Veh], 1 μM and 10 μM). (C) Images and graphs show PCR bands and relative mRNA levels of *Hccll11* at different time points (Veh, 1, 3, and 6 h) under 10 μM NE stimuli. (D) The graph shows ELISA analysis of eotaxin-1 protein level in the CCD-18co cell culture serum at different time points (Veh, 1, 3, and 6 h) under 10 μM NE stimuli. Data were collected from four independent experiments in (B), (C) and (D), respectively. (E) Graphs show the gating strategy of colonic fibroblasts using FACS. (F) Images show PCR bands of adrenoceptors expressed in sorted fibroblasts from the colon of the human CCD-18co cell line (top) and the mice (bottom). Data are represented as mean ± SEM. One-way ANOVA was performed and compared with the Veh groups. **P < 0.01.

To examine whether colonic fibroblasts express ARs, we sorted mice colonic cells using flow cytometry. After excluding CD45^+^ and CD31^+^ cells, we identified CD34^+^ PDGFRα^+^ cells as fibroblasts based on previous research ^36^ (Figure 5E). PCR analysis of the sorted fibroblasts revealed the expression of α1B and β2 ARs, consistent with the expression pattern observed in human CCD-18co cells (Figure 5F). These findings suggest that intestinal fibroblasts respond to NE stimulation by releasing eotaxin-1, thereby inducing eosinophilic infiltration into the colon.

### Eotaxin-1 is essential for sympathetic activity-induced eosinophil infiltration and colorectal hypersensitivity in the MS model

After demonstrating that NE signalling modulates eosinophil infiltration via a mesenchymal cell-dependent mechanism in the colon, we subsequently examined whether the same underlying mechanism applies to the MS-induced IBS model. Eotaxin-1-ir cells were significantly increased in the colonic mucosa of MS rats (Figure 6A) but were reduced after 6-OHDA treatment (Figure 6A), supporting the role of sympathetic signalling. Additionally, 6-OHDA treatment suppressed visceral hypersensitivity in MS rats, as evidenced by a reduced VMR at 60 mmHg to CRD (Figure 6B).

**Figure 6.**
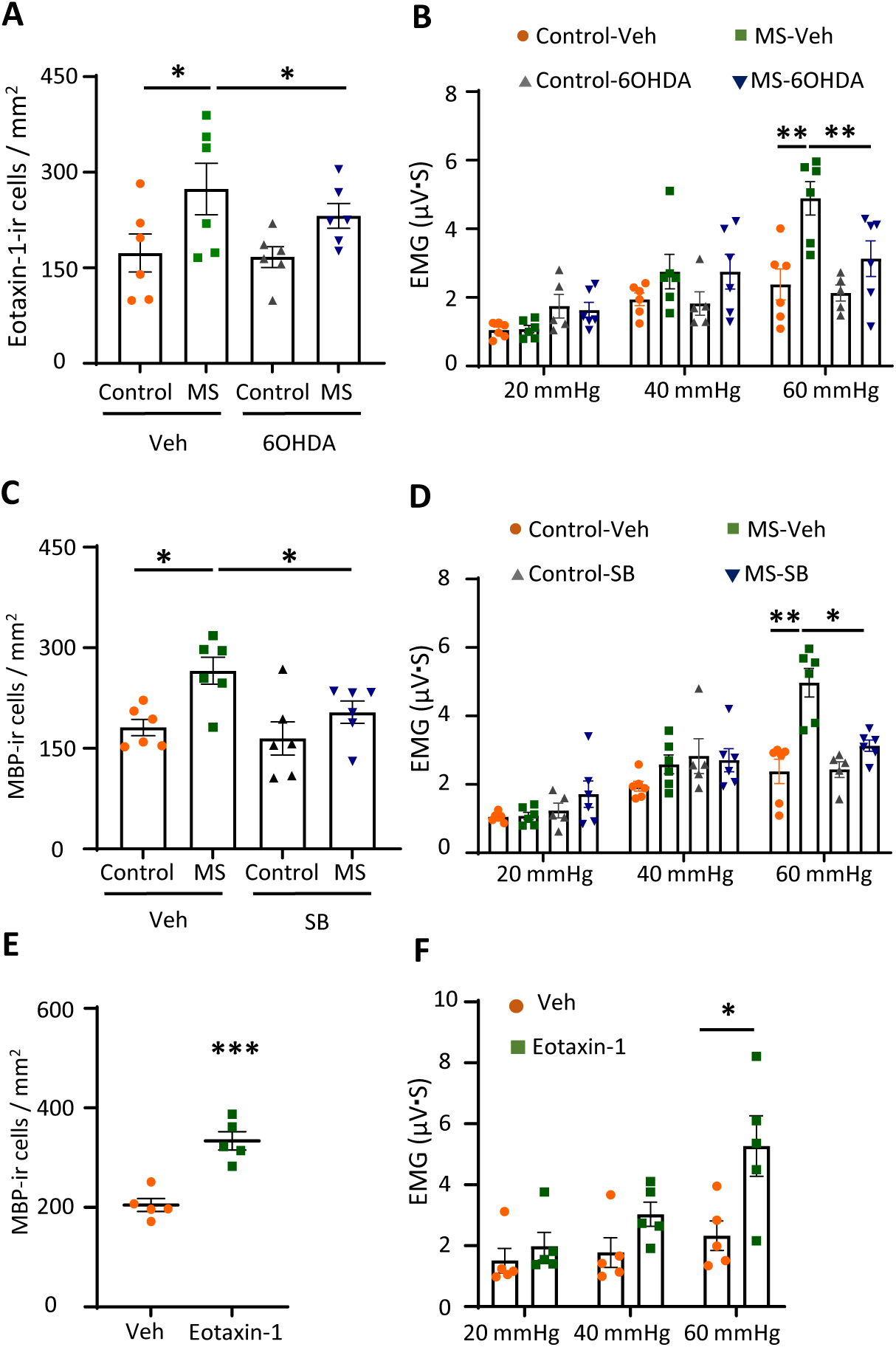
Eotaxin-1 is essential for eosinophil infiltration and colorectal hypersensitivity in the maternal separation (MS) model. (A) Bar graphs show numbers of eotaxin-1-ir cells in control and MS rats with/without 6-OHDA treatment (n= 6 in each group). (B) Bar graphs represent the data of electromyography (EMG) response to colorectal distention (CRD) in control and MS rats with/without 6-OHDA treatment (n = 6 in each group). (C) The bar graph shows the numbers of major basic protein-immunoreactive (MBP-ir) cells in the control and MS groups with/without SB328437 treatment (n = 6 in each group). (D) Bar graphs represent data of EMG response to CRD in control and MS rats with vehicle (Veh)/SB328437 treatment (n = 6 in each group). (E) The graph shows the number of MBP-ir cells following eotaxin-1 treatment (n = 5 in each group). (F) Bar graphs represent EMG response to CRD following Veh/eotaxin-1 (n = 5 in each group). Data are represented as mean ± SEM. Two-way ANOVA with Bonferroni post hoc test was used for (A-D and F), and an unpaired *t-*test was used for (E). *P < 0.05, ***P < 0.001.

As eotaxin-1 is the main chemoattractant causing eosinophil infiltration, we administered the eotaxin-1 receptor CCR3 antagonist SB328437 for 7 continuous days to both control and the MS animals and measured eosinophil infiltration and visceral response to CRD. Treatment with SB328437 significantly suppressed both visceral hypersensitivity and eosinophil infiltration in MS animals (Figure 6C, D).

To investigate whether colonic adrenaline signalling can induce visceral hypersensitivity and determine the involvement of eotaxin-1, we measured the VMR to CRD 6 h after colonic NE infusion. Colorectal NE infusion induced colorectal hypersensitivity at 60 mmHg (Supplemental Figure S4C), which was inhibited by the eotaxin-1 receptor CCR3 antagonist SB328437 (Supplemental Figure S4D).

We also administered an eotaxin-1 colorectal infusion to directly evaluate its effect on eosinophil infiltration and visceral hypersensitivity. Eotaxin-1 infusion induced significant eosinophil infiltration into the colorectal mucosa (Figure 6E). Rats that received eotaxin-1 colorectal infusion exhibited hypersensitivity to CRD (Figure 6F).

Collectively, these data suggest that overactive sympathetic signalling results in eotaxin-1 release, ultimately inducing eosinophil infiltration and visceral hypersensitivity.

### Upregulation of MBP- and eotaxin-1-ir cells in the colorectal tissue of patients with IBS

Previous clinical studies suggest that a subset of patients with IBS show eosinophil infiltration in their GI tract ^37, 38^. To ascertain whether the presence of eotaxin-1-expressing cells, as observed in animal models, could also be identified in humans with IBS, biopsy specimens of colorectal tissue were obtained from both healthy controls and patients with IBS. We observed a pronounced elevation in the number of MBP-ir cells in colorectal tissues of patients with IBS, accompanied by a tendency for an increased presence of tryptase-ir cells (Figure 7A-D). Given the pivotal role of eotaxin-1 released from mesenchymal cells in eosinophil infiltration, we conducted double staining for eotaxin-1 and vimentin. Increased eotaxin-1-ir cells were detected in the colorectal tissues of patients with IBS, accompanied by an augmentation in the number of cells co-expressing vimentin, compared with those in the healthy controls (Figure 7E, F). Consequently, eotaxin-1 in vimentin-positive cells (which are likely to be fibroblasts) plays a pivotal role in the onset of microinflammation, which is a key factor in the pathogenesis of IBS.

**Figure 7.**
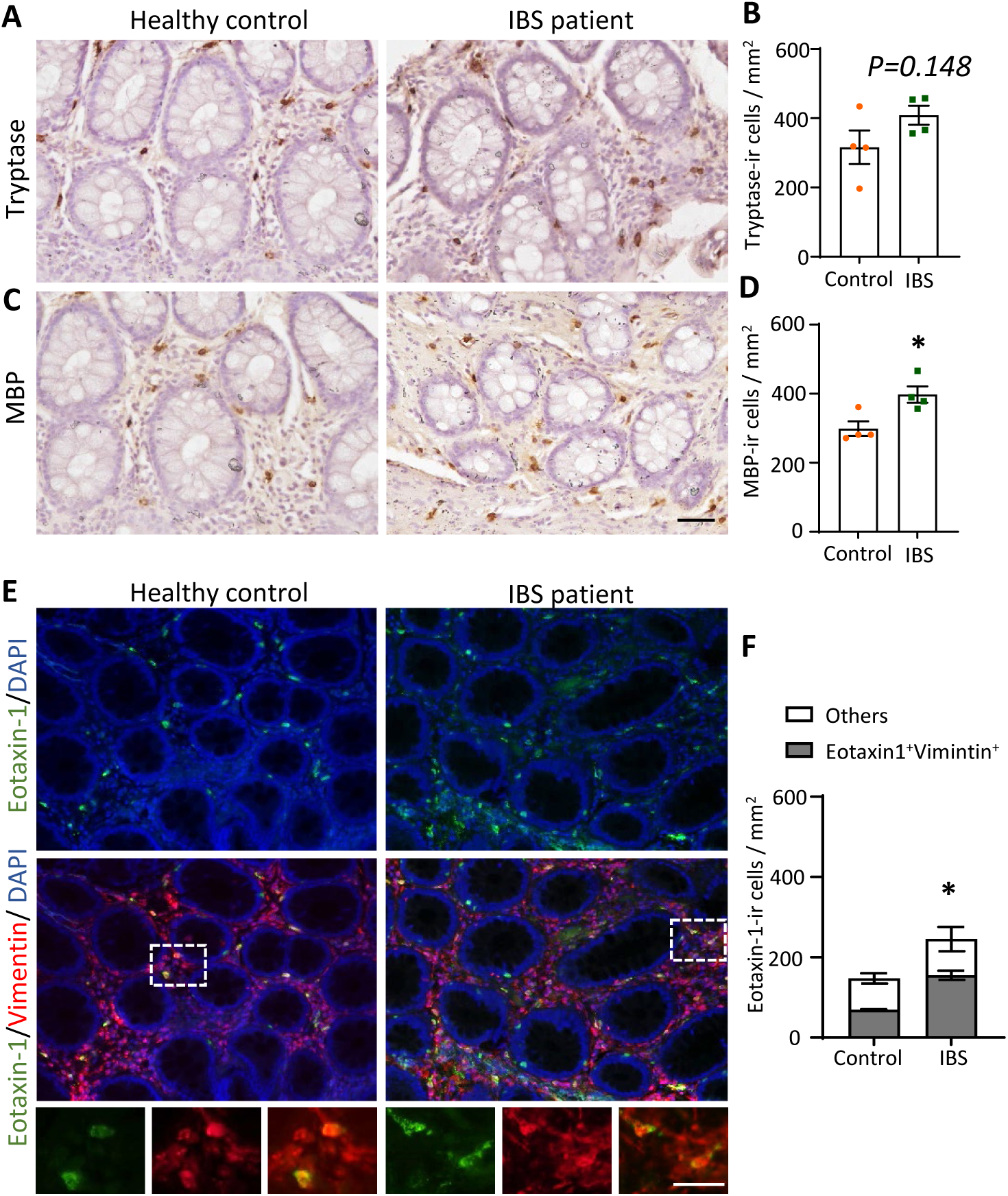
Upregulation of major basic protein (MBP)- and eotaxin-1-immunoreactive (ir) cells in colorectal tissues of patients with irritable bowel syndrome (IBS). (A-B) Representative images of tryptase (A) and the number of tryptase-ir cells (B) in healthy controls and patients with IBS. (C-D) Representative images of MBP (C) and the number of MBP-ir cells (D) in healthy controls and patients with IBS. (E) Representative images of eotaxin-1 and vimentin in healthy controls and patients with IBS. (F) Number of eotaxin-1-ir positive cells and Eotaxin-1^+^Vimintin^+^ cells in controls and patients with IBS (n = 4 in each group). Data are presented as mean ± SEM, with statistical analysis performed using the nonparametric Mann-Whitney test. *P < 0.05.

## DISCUSSION

In the present study, we employed a rat model of MS stress and successfully reproduced the main IBS characteristics, including microinflammation and visceral hypersensitivity, confirming that MS rats displayed sympathetic overactivity. 6-OHDA-induced sympathetic nerve degeneration reduced eosinophil-associated microinflammation in MS rats. Using chemogenetic approaches, we demonstrated that sympathetic activation induces eosinophil infiltration and visceral hypersensitivity in a mesenchymal cell-dependent manner in the colons of mice. Furthermore, patients with IBS showed upregulation of mesenchymal cells-derived eotaxin-1, associated with eosinophil infiltration in their colorectal tissue. These findings reveal that the SNS overactivation-induced release of mesenchymal cell-derived eotaxin-1 could be a potential inducer of eosinophil-associated mucosal microinflammation in patients with IBS.

Patients with IBS exhibit immune alterations in the GI ^5^. Mast cells have been well-studied in both clinical and experimental animal research and implicated as important contributors to IBS-associated microinflammation. However, reports on mucosal mast cell counts are controversial, with some studies reporting upregulated, unchanged, or downregulated counts in patients with IBS ^6^. Notably, a significant subset of patients with IBS does not exhibit mast cell alterations in the GI mucosa ^7, 8, 39–41^, suggesting the involvement of additional mechanisms underlying microinflammation in IBS. Herein, no significant alterations were observed in the number of mast cells in the mucosal tissue in both the MS model and the sympathetic activation condition despite a discernible trend towards a significant increase. Intestinal mast cells respond to various allergens, pathogens, and other agents that can be ingested, inhaled, or encountered after intestinal lumen epithelial barrier disruption ^42^. The subtle alterations in mast cells may be attributed to slight changes in the luminal environment in MS rats.

Eosinophils are innate immune cells that play a pivotal role in the GI and maintain intestinal homeostasis ^32^. Emerging clinical studies have reported that a subpopulation of patients with IBS exhibits colonic eosinophil infiltration and activation ^4, 5, 9, 10, 37, 38^. Our study findings, showing eosinophil-dominant microinflammation, may reflect these clinical manifestations.

Eosinophil proliferation, maturation in the bone marrow, and migration to the circulation are regulated by IL5 released from ILC2 cells ^33, 43^. The recruitment of eosinophils to the GI is regulated primarily by eotaxin-1. We observed that IL5 expression did not change in both the bone marrow and mucosa, whereas expression of the eotaxin-1 gene increased in the intestinal mucosa after NE stimulation or in MS rats. This result suggests that sympathetic activation induces eosinophil infiltration via the eotaxin-1 pathway rather than the ILC2 pathway.

In healthy individuals, eotaxin-1 is responsible for the normal recruitment of eosinophils to the GI and is continuously expressed in the intestinal lamina propria ^34^. Eotaxin-1 deficient mice exhibit a large, selective reduction in eosinophils residing in the intestinal tract ^34^. Eotaxin-1 is produced by various cell types, including macrophages, fibroblasts, and epithelial cells ^44^. We found that in the intestinal mucosa, eotaxin-1 was mainly expressed by macrophages and vimentin-positive mesenchymal cells but not by endothelial cells. After NE stimulation, the number of mucosal eotaxin-1-expressing mesenchymal cells (but not macrophages) increased, indicating that mesenchymal cells are the main source of eotaxin-1 in response to sympathetic signalling. These vimentin-positive mesenchymal cells in the intestinal mucosa were defined as lamina propria fibroblasts or myofibroblasts. Moreover, both the human fibroblast cell line and mice colonic fibroblasts expressed α1B and β2 ARs (Figure 5F). These data suggest that intestinal fibroblasts are regulated by sympathetic signalling and play a crucial role in eosinophil migration and infiltration through the release of eotaxin-1. In our study, both findings from animal models and human samples suggest that fibroblasts releasing eotaxin-1 played a role in the microinflammation via sympathetic signalling rather than antigen triggers. Our findings highlight a novel mechanism through which fibroblasts participate in IBS pathogenesis and suggest potential therapeutic targets for this complex disorder.

On the other hand, fibroblasts constitute a crucial component of the GI stroma and substantially contribute to wound healing, extracellular matrix remodelling, and immune responses. In inflammatory conditions such as IBD, fibroblasts act as major sources of inflammatory cytokines and chemokines, perpetuating chronic tissue inflammation ^45, 46^. The current observations suggest that fibroblasts can be regulated by adrenergic signalling, thereby offering a novel underlying mechanism and potential approach for treating these diseases or conditions.

Visceral pain is a key pathological feature of IBS, which involves the central and peripheral nervous systems ^47, 48^. Herein, we demonstrated that eosinophil infiltration contributed to colorectal hypersensitivity in an MS stress-induced IBS model. Nevertheless, we did not focus on the mechanisms through which eosinophil infiltration leads to visceral hypersensitivity, a phenomenon that has been well studied and documented in previous studies ^25, 49^. Eosinophils can release various bioactive mediators, such as MBP, EPO, cationic proteins, neuropeptides, and reactive oxygen species. These mediators may affect receptors in the peripheral afferent, resulting in visceral hypersensitivity ^25, 50^.

Our study focused on investigating the role of the overactive peripheral SNS on immune alteration using an IBS animal model. The results suggest a potential mechanism underlying microinflammation and visceral hypersensitivity. Nevertheless, we acknowledge the contribution of a dysregulated HPA axis in the IBS pathogenesis and the contribution of mast cells to microinflammation in IBS. We believe both mast cells and eosinophils contribute to intestinal microinflammation in the complex pathogenesis of IBS.

## Conclusion

The present study revealed that SNS overactivation led to eosinophil infiltration in the colon of the MS stress-induced IBS model. This infiltration can be attributed to the release of eotaxin-1 from mesenchymal cells, contributing to the development of visceral hypersensitivity. Our findings not only suggest that fibroblast-derived eotaxin-1 and eosinophil could be potential targets for therapeutic interventions in IBS but also provide evidence for understanding the effectiveness of psychotherapy, which can relieve sympathetic activity in IBS management.

## Contributors

YD conceived and designed research; SD conducted the experiments and drafted the manuscript; HK, KN, and YD supervised the study; FZ performed a part of animal experiments; TK prepared the AAV for experiment of DBH-cre mice; YO and TT contributed to FACS experiments; YH and KM contributed to the experiments related to Cartpt-cre mice; HF and SS conducted the clinical study and provided the biopsy specimens; SD and MO performed the double-blind data analyses for clinical data. HK, YC, KN, and YD edited and revised the manuscript. All contributing authors have agreed to the submission of this manuscript for publication.

## Acknowledgements

We thank Prof. Kazuto Kobayashi for providing DBH-cre mice and Seiko Oki, Serika Yamada, Mitsue Hagihara, and Mayumi Yamada for providing technical support during this study and preparation of the manuscript.

## Ethical approval

Animal experiments were approved by the Committee on Animal Research at Hyogo Medical University (No. 2021-22, 2021-18, 2021-04). Human studies were performed in accordance with the Declaration of Helsinki and approved by the Ethics Review Board of Hyogo Medical University (No. 4706).

## Patient consent

All patients and healthy controls provided written informed consent.

## Data availability statement

Data are available upon reasonable request.

