## supplemental for "Sympathetic Nervous System Overactivation Induces Colonic Eosinophil-Associated Microinflammation and Contributes to the Pathogenesis of Irritable Bowel Syndrome"

### SUPPLEMENTARY INFORMATION

#### MATERIALS AND METHODS

##### Animals and protocols

The animals in the study were housed and fed under the exact same conditions. Following the 3R principles, the experimental protocols were designed to minimize the number of animals used. The number of animals used in each group is mentioned separately in the figure legends. A double-blind method was employed in experiments of DBH-cre mice and immunohistochemistry of clinical biopsy specimens. A researcher unrelated to the experiment randomly allocated the animals to each group. The grouping was revealed only after the experiments. Data were analysed by the first author.

##### Electrode implantation

Electrodes were implanted into the external oblique muscles of the rats to record muscle contraction, as reported previously [1]. Surgery was performed under 2% isoflurane-induced general anaesthesia. Two wires (Cooner Wire, Chatsworth, CA, USA) were implanted into the left external oblique muscle, and one wire was implanted on the other side. Wires were externalised at the backs of the animals, and the skin was closed with a suture. The animals were allowed 5 days to recover before the next procedure.

**Colorectal distention and electromyography recording**

The control and MS rat models were trained to acclimate to a cylindrical box (length: 15 cm; diameter: 5 cm) to perform colorectal distention (CRD) calmly. CRD was performed as described previously [1]. A 4 cm long slack latex balloon (Okamoto, Tokyo, Japan) attached to a 10-cm long gavage needle was inserted into the colon through the anus. Varying pressure (20, 40, and 60 mmHg) was applied to the colon by inflating the balloon with 37 °C water. Each pressure was tested for 20 s at a 5 min interval time. The visceromotor response (VMR) to CRD was recorded using electromyography (EMG) with wires implanted in the muscle. The EMG signals were filtered at 1000 Hz and recorded using Power Lab (AD Instruments, Castle Hill, New South Wales, Australia). The area under the curve (AUC) was calculated for 10 s at baseline and during CRD. The VMR at each CRD pressure was normalised to the baseline. The threshold was calculated from a ramp distention experiment independently, and the pressure of the CRD at which the VMR increased to five times the baseline value was characterised as the threshold.

**Adeno-associated virus**

The plasmid of capsid AAV PHP.S, pUCmini-iCAP-PHP.S (Addgene, Plasmid No. 103006), AAV Helper plasmid (Takara, Tokyo, Japan), and shuttle plasmid, and AAV-hSyn-DIO-hM3D(Gq)-mCherry (Addgene, Plasmid No. 44361) were transfected in HEK293T cells (RIKEN BRC, Japan). Virus particles were purified via OptiPrep ultracentrifugation, concentrated using Amicon Ultra-15 centrifugal filter unit (Millipore, Bedford, MA), and kept in 1× PBS. Titre was measured via quantitative polymerase chain reaction with SYBR Green (Takara).

AAV8 hSyn-DIO-hM3D (Gq)-mCherry (Addgene #44361) and AAV8 hSyn-DIO-mCherry (Addgene #50459) vectors were obtained from Addgene.

**AAV injection**

*Retro-orbital injection:* Four-week-old DBH-Cre mice were anaesthetised with 2% isoflurane (Wako Pure Chemical Corporation, Osaka, Japan). The mouse was positioned on its side, and the loose skin around the eyes was pulled back using the fingers. The eye protruded slightly, and a 30-G needle of an insulin syringe (BD, New Jersey, USA) was inserted at an angle of approximately 45 ºC. The needle was passed through the inferior conjunctiva into the retroorbital sinus [2]. A 150 μL dose of AAV ($1 \times 10^{13}$ viral genomes/mL) was injected into the sinus, and the needle was gently removed to prevent eye damage. The next procedure was performed 4 weeks after AAV injection.

*Spinal cord injection:* In the experiment reported in figure 2I, an incision was made around the thoracic region of the spinal cord in cartpt-cre mice. To expose the spinal cord, the paraspinal muscles and dorsal vertebral elements were carefully excised using fine forceps. A glass pipette prepared for a microinjector (#UMP3; WPI) was utilized to administer 100 nL of the AAV vector with 0.1% Fast Green (#F7252, Sigma-Aldrich) at 10 distinct sites (infusion rate: 60–70 nL/min) bilaterally.

**Colorectal infusion**

The rats or mice used for colorectal infusion experiments were anaesthetised with 2% isoflurane, and a plastic oral gavage needle was slowly inserted into the colon through the anus. The needle was inserted 2 cm from the anus, and NE or saline was injected slowly to avoid leakage. After the injection, the animals were allowed to keep their hips elevated for 10 min. Norepinephrine (Sigma-Aldrich, Burlington, MA, USA) was injected at 0.5 mg/kg. The next steps were performed 6 h after NE. Eotaxin-1 (PeproTech, New Jersey, USA) was injected at 1 mg/kg for rats, and EMG was recorded 24 h after the injection.

**Colon lamina propria cell harvesting and flow cytometry**

Colon tissues of wild-type mice 6 h after NE colorectal infusion and those of DBH-cre mice after 5 days of continuous clozapine *N*-oxide / saline stimulation were collected under 2% isoflurane anaesthesia. Samples were placed in cold Hank's balanced salt solution (HBSS), and faecal pellets, blood vessels, and mesenteric tissue were carefully removed. The samples were cut along the longitudinal axis into 1 cm pieces and placed in HBSS containing 1 mM dithiothreitol-ethylenediaminetetraacetic acid and 10% foetal bovine serum (FBS). The samples were shaken at 37 °C and 150 rpm for 20 min to remove the epithelial cells. The samples were then placed in Roswell Park Memorial Institute 1640 medium (Nacalai, Kyoto, Japan), containing 10% FBS, 1 mg/mL Collagenase IV (Sigma-Aldrich, Burlington, MA, USA), and 1 mg/mL Dispase II, and shaken for 60 min to obtain a single-cell solution. The single-cell solution was centrifuged through discontinuous Percoll (Cytiva, Tokyo, Japan) gradients (40%: 80%) to collect cells from the lamina propria. The collected cells were washed with HBSS, and the cell density was adjusted to $1 \times 10^6$ cells/mL.

The cells were resuspended in HBSS containing 10% FBS and incubated with fluorescently conjugated antibodies for 30 min on ice. The following fluorescently conjugated antibodies were purchased from BioLegend and used: PE/Cy7 anti-mouse CD45 (30-F11, 1:500), FITC anti-mouse CD11b (M1/70, 1:200), PE anti-mouse Siglec-F (S17007L, 1:200), PerCP/Cy5.5 anti-mouse FceRIa (MAR-1, 1:200), APC anti-mouse Ly-6G (1A8, 1:200), BV711 anti-mouse F4/80 (BM8, 1:200), BV711 anti-mouse CD117 (2B8, 1:200), FITC anti-mouse CD3 (145-2C11, 1:200), PE anti-mouse CD4 (GK1.5, 1:200), and APC anti-mouse CD8 (53-6.7, 1:200); 4',6-diamidino-2-phenylindole (DAPI, 1:1000) was also purchased. After washing the cells with PBS containing 0.2% BSA, the fluorescence intensity of individual cells was determined using a FACSAriaIIIu flow cytometer (BD Bioscience, Franklin Lakes, New Jersey, USA). Dead cells were stained with DAPI and excluded from the analysis. For analysis, the adhesion signals were excluded by FSC-A(area) and FSC-H(height) at first, and then approximately 50,000 events were gated to live cells using DAPI in each sample. Subsequently, the gates were set on anti-CD45 positive events, followed by immune cell gating separately. FlowJo software (BD Bioscienes, Ashland, OR, USA) Version 10.8.1 was used for data analysis.

**Sorting of CD34+ PDGFRα+ cells**

A single-cell suspension from the colon lamina propria was prepared as described above. Cells were resuspended in Hank's Balanced Salt Solution (HBSS) containing 10% foetal bovine serum (FBS) and incubated with fluorescently conjugated antibodies for 30 min on ice. The following fluorescently conjugated antibodies were purchased from BioLegend (San Diego, CA, USA) and used: APC anti- mouse CD34 (MEC14.7, 1:200), PE anti-mouse PDGFRα (APA5, 1:200), FITC anti-mouse CD31 (390, 1:200), and PE/Cy7 anti- mouse CD45 (30-F11, 1:500). After washing and resuspending in HBSS containing 2% FBS, mouse fibroblasts (CD45- CD31- CD34+ PDGFRα +) were sorted via BD FACSAriaIIIu (BD Biosciences). Endothelial and immune cells were excluded as CD31- and CD45-positive cells, respectively.

**Immunohistochemistry**

Immunohistochemistry (IHC) was performed as described previously [1]. Briefly, rat and mice were anaesthetised with 2% isoflurane and transcardially perfused with 1% paraformaldehyde, followed by 4% paraformaldehyde. The brain, superior cervical ganglion, DRG, spinal cord, and gastrointestinal tract (rat: 3 – 5 cm from anus; mouse: all colon tissue) tissues were dissected and post-fixed in 4% paraformaldehyde for 24 h at 4 °C. The tissues were then transferred to 30% sucrose for 2 days and finally embedded in Tissue-Tek O.C.T. (Sakura Finetek). In a parallel experiment, fresh tissues were dissected without paraformaldehyde perfusion and embedded in Tissue-Tek OCT. Frozen samples were sectioned at 10 μm thickness using a cryostat (Thermo Fisher Scientific, MA, USA) and prepared for IHC staining. Fresh sections were post-fixed using acetone. Human paraffin sections were first incubated with 0.4% pepsin for 20 min at 37 °C and then with primary antibodies overnight at 4 °C, followed by secondary antibody incubation for 1 h at room temperature. The primary and secondary antibodies used in this study are listed in Tables 1 and 2. As reported previously, signals were obtained via immunofluorescence or 3, 3′-diaminobenzidine (DAB) staining [1]. Sections were observed under a microscope (Nikon, Eclipse 80i, Nikon Instruments, Japan), and images were taken at 10× or 20× magnification using NIS-Elements D 3.2 software. Three sections were included for each animal, and 4 – 6 animals were included in each group. Cell counts were analysed using the ImageJ and Fiji software.

**H&E staining**

10 μm thick sections of colon tissue were prepared as described above. The sections were stained with Mayer's Haematoxylin (Sigma, St. Louis, USA) for 2 min followed by Eosin (sigma St. Louis, USA) for 15 sec. Subsequently, the sections were dehydrated and sealed.

**Double staining**

To exclude nonspecific immunofluorescence signals in the colonic samples, we performed double IHC of certain antibodies with DAB combined with alkaline phosphatase (AP) staining. Incubation with primary antibodies was performed as described above. The sections were then incubated with horseradish peroxidase-conjugated or AP-conjugated secondary antibodies separately for 1 h at room temperature. Horseradish peroxidase was detected using DAB staining as previously described [1]. AP was detected using the ImmPACT Vetor Red AP Substrate Kit (SK-5105; Vector, Newark, CA, USA) according to the manufacturer's instructions.

**Mouse-on-mouse staining**

We used a mouse anti-cre antibody from Sigma-Aldrich to detect DBH-cre, MBP, and eotaxin-1 positive cells. MOM immunodetection kit (BMK-2202; Vector, Newark, CA, USA) was used for staining according to the manufacturer's instructions.

**Enzyme-linked immunosorbent (ELISA) assay**

*Eotaxin-1 ELISA:* CCD-18Co cells were cultured in 6-well plates at a density of $5 \times 10^4$ /mL and incubated at 37 °C under 5% $CO_2$ before NE stimulation. The medium was replaced with 0, 1, and 10 μM NE and incubated for 1, 3, or 6 h. After incubation, the supernatants were harvested for eotaxin-1 ELISA assay using the Human CCL11/Eotaxin Kit (R&D Systems, Inc. Canada) according to the manufacturer's protocol.

*NE ELISA:* Blood samples were collected from the veins of adult MS rats or DBH-cre mice after 5 days of continuous clozapine *N*-oxide/saline stimulation under 2% isoflurane anaesthesia and kept for 6 h at 4 °C to clot. The clot was removed by centrifugation at 1,500 *g* and 4 °C for 10 min. The supernatants were subjected to NE ELISA (LifeSpan Biosciences. Inc., Seattle, WA, USA) according to the manufacturer's instructions.

**Reverse transcription-polymerase chain reaction**

*CCD-18Co cells:* CCD-18Co cells (American Type Culture Collection, VA, USA) were cultured in 6-well plates as described above. The cells were stimulated with 0, 1, and 10 μM NE and incubated for 1, 3, or 6 h. After incubation, the cells were collected for polymerase chain reaction (PCR) analysis. Total RNA was extracted using TRIzol (Sigma-Aldrich, Burlington, MA, USA), and PCR amplification of *ccl11* (eotaxin-1) was performed as described previously [1].

*Sorted cells:* $CD34^{+}$ $PDGFR\alpha^{+}$ sorted by FACS were collected in a tube; total RNA extraction and PCR protocols were the same as described above.

*Colonic tissue:* Six hours after NE colorectal infusion, rats were anaesthetised with 2% isoflurane. The colon tissue was dissected and rapidly frozen on dry ice. The colonic tissues were dissected from control and MS model rats at 7 – 8 weeks old and frozen on dry ice. Total RNA was extracted, and PCR was performed as above.

*Bone marrow*: bone marrow was dissected from the hind limb long bones (femurs) from the control or MS rat after sacrifice. Total RNA was extracted, and PCR was performed as above.

The primers used in the reverse transcription-PCR are listed in Table 3.

**Eosinophil Peroxidase (EPO) Assay**

EPO activity was tested as our previous reports [1]. 0.5 cm-long colon tissues were taken from rats after sacrifice. For the EPO assay, tissue samples were homogenized in 50 mM Tris·HCl (pH 8.0) with 0.1% Triton X-100 at a ratio of 1 mL per 50 – 100 mg of tissue. The homogenate was centrifuged at 1,600 g for 5 minutes at 4°C. The resulting supernatants were used to measure EPO activity, utilizing the oxidation of o-phenylenediamine (OPD) in the presence of H2O2. For the assay, 20 μL of the supernatant was combined with 580 μL of 50 mM Tris·HCl buffer containing 10 mM OPD (Nacalai,

Kyoto, Japan) and 4 mM $H_2O_2$. Enzyme activity was determined photometrically at 492 nm following a 5 min incubation.

**Faecal pellets analysis**

After intraperitoneal injection of Clozapine *N*-oxide (CNO) or vehicle, DBH-cre mice were placed in a clean cage for 2 h for faecal pellet collection. The number of faecal pellets in the clean cage was counted, and the data were recorded once a day for 5 days. Data are presented as the number of pellets formed per hour.

**6-OHDA treatment**

To degenerate SNS activity, 6-OHDA (Cayman Chemical, Ann Arbor, MI, USA) was dissolved in saline containing 1% ascorbic acid. Seven-week-old rats (MS and control) were intraperitoneally administered with 6-OHDA (150 mg/kg). The vehicle group was administered the same solution but without 6-OHDA. Subsequent experiments were performed 7 days after 6-OHDA treatment.

**Pharmacological agents**

Clozapine *N*-oxide (Cayman Chemical, Ann Arbor, Michigan, USA) was dissolved in saline containing 0.5% dimethylsulphoxide (DMSO) and administered intraperitoneally (1 mg/kg) for 5 continuous days. The eotaxin-1 CCR3 receptor antagonist SB328437 (Tocris Bioscience, Bristol, United Kingdom) was intraperitoneally injected at a dose of 1 mg/kg with 10% DMSO and 0.1% Tween 20. The pan-alpha adrenergic receptor (AR) antagonist phentolamine (R&D Systems, Minneapolis, USA; 1 mg/kg in saline) and the pan-beta AR antagonist propranolol (pro) (R&D Systems, Minneapolis, USA; 10 mg/kg in saline) were administered intraperitoneally. Before experiments, the MS model was treated with SB328743, phentolamine, and propranolol for 7 continuous days and 1 h before the NE colorectal infusion. The treatment of each group was finished within short time intervals to minimise potential confounders.

Table 1: Primary antibodies used for immunohistochemical staining

| Antigen | Host | Manufacture | Cat. number | Dilution |
|---|---|---|---|---|
| Cre Recombinase | Mouse | Sigma-Aldrich | MAB3120 | 1:500 |
| RFP | Rabbit | MBL | PM005 | 1:1000 |
| eotaxin-1 | Mouse | Santa Cruz | Sc-373767 | 1:50 |
| MBP | Mouse | Bio-Rad | MCA5751 | 1:500 |
| tryptase | Mouse | Abcam | ab2378 | 1:500 |
| MPO | Rabbit | Abcam | Ab9535 | 1:1000 |
| Iba1 | Rabbit | Wako | 019-1074 | 1:1000 |
| CD68 | Rabbit | Abcam | Ab2512 | 1:1000 |
| CD31 | Mouse | Millipore | MAB1393 | 1:1000 |
| vimentin | Rabbit | Abcam | Ab92547 | 1:5000 |

Table 2: Secondary antibodies used for immunohistochemical staining

| Antibody | Manufacture | Cat. number | Dilution |
|---|---|---|---|
| Biotinylated-horse anti- mouse IgG | Vector | BA2001 | 1:200 |
| Biotinylated-goat anti-rabbit IgG | Vector | BA1000 | 1:200 |
| Donkey anti-rabbit Alexa Fluor 594 | Molecular Probes | A32754 | 1:1000 |
| Goat anti-rabbit Alexa Fluor 594 | Molecular Probes | A11037 | 1:1000 |
| Goat anti-mouse Alexa Fluor 488 | Molecular Probes | A11029 | 1:1000 |
| Donkey anti-goat Alexa Fluor 488 | Molecular Probes | A11055 | 1:1000 |

Table 3: Sequences of primers used in PCR

| Gene | Accession No. | Type | Sequence (5′–3′) |
| --- | --- | --- | --- |
| *hccl11* | NM_002986.3 | sense | AAGCTCACACCTTCAGCCTC |
| | | antisense | CACTCAGGCTCTGGTTTGGT |
| *rccl11* | NM_019205.2 | sense | CAGCTCTCCACAGCACTTCT |
| | | antisense | GGGTGCCGATATTCTCCCAT |
| *rccl24* | NM_001013045.1 | sense | CTTGCACCCCAGCTTTGAAC |
| | | antisense | GGTGCTATTGCCTCGGAGTT |
| *rccl26* | XM_006249188.4 | sense | GGTTCTTGAGCGTCCACACA |
| | | antisense | TGGCTGGACACAGTATTGCT |
| *rIL5* | NM_021834.1 | sense | GATGCTTCTGTGCTTGAGCG |
| | | antisense | TCTTCGCCACACTGCTCTTT |
| *rIL33* | NM_001014166.1 | sense | CTGCACAATCAGGAGACGGT |
| | | antisense | CCCAGAAGGCACAGACCTTT |
| *rPrg3* | XM_031373412.1 | sense | GCAGGGGAGAGAGTTGGTTC |
| | | antisense | GACGCCTCCAATCCAGACAA |
| *rTpsab1* | NM_019322.2 | sense | TGACTTCTACATCGCCCAGG |
| | | antisense | Prg-GGCAGGAGTCATGTCCTTCA |
| *mAdrb1* | NM_007419.3 | sense | TCTGGTCATGGGATTGCTGG |
| | | antisense | CCTGTTGGTGACGAAATCGC |
| *mAdrb2* | NM_007420.3 | sense | GAACGACAGCGACTTCTTGC |
| | | antisense | GATCCACTGCAATCACGCAC |
| *mAdrb3* | NM_013462.3 | sense | GAGTGAGTCCCCTGGAACCT |
| | | antisense | AGTGAGGAGACAGGGATGAAA |
| *mAdra1a* | NM_001271760.1 | sense | CAACCCGAGCTGCAAAGTTC |
| | | antisense | GGAAACGTGAGCCTGAGGAA |
| *mAdra1b* | XM_011248675.4 | sense | ATTGAAAGCAGACCCTCCTCG |
| | | antisense | GCAGGTGCTGATGTGTTGTG |
| *mAdra1d* | NM_013460.5 | sense | GCCACTCGCTCAAGTATCCA |
| | | antisense | GACGATGGCTAGGGTCTTGG |
| *mAdra2a* | NM_007417.5 | sense | CGCTGGACCTAGAGGAGAGT |
| | | antisense | TTCAGCGAGCTGTTGCAGTA |
| *mAdra2b* | NM_009633.4 | sense | CCAACAGTAGCGGAGCTAGG |
| | | antisense | TGTCCAGACTGATGGCACAC |
| *mAdra2c* | NM_007418.3 | sense | TCACCGTGGTAGGCAATGTG |
| | | antisense | GCGGTAGAACGAGACGAGAG |
| *mCD45* | NM_001268286.1 | sense | CCAGTGATGCTACCACAACGA |
| | | antisense | GCACGAAGGTTGTCCAACTG |
| *mCD31* | NM_001032378.2 | sense | GTGAATGACACCCAAGCGTT |
| | | antisense | GAGCCTTCCGTTCTCTTGGT |
| *mCD34* | NM_001111059.2 | sense | ACAGTACCTCACAACCCTGC |
| | | antisense | GTCCAGGGCAAGTGCTACAT |
| *m Pdgfra* | NR_144636.1 | sense | CACTTTGACCGTCCCCAAGG |
| | | antisense | CATCCCGACCACACAAGAACA |
| *hAdrb1* | NM_000684.3 | sense | ACTCGAAGCCCACAATCCTC |
| | | antisense | TCTGGCTGGTAGTGTGTTCC |
| *hAdrb2* | NM_000024.6 | sense | CTCTCATCGTCCTGGCCATC |

| | | | |
|---|---|---|---|
| | | antisense | GAATGATCACCCGGGCCTTA |
| *hAdrb3* | NM_000025.3 | sense | GCCAATTCTGCCTTCAACCC |
| | | antisense | TCGTCAGGTTCTGGAGGGTA |
| *hAdra1a* | NM_001322502.1 | sense | AATGCTTCCGACAGCTCCAA |
| | | antisense | GATGATGCAGAGGCCCATGA |
| *hAdra1b* | NM_000679.4 | sense | CTGTTGAGCTTCACCGTCCT |
| | | antisense | ATCCTCAGGGTCAGCTCCTT |
| *hAdra1d* | NM_000678.4 | sense | ACTCACTCAAGTACCCAGCC |
| | | antisense | CTCACGGGAGAACTTGAGCA |
| *hAdra2a* | NM_000681.4 | sense | GGCTACTGGTACTTCGGCAA |
| | | antisense | CTGGTAGATGCGCACGTAGA |
| *hAdra2b* | NM_000682.7 | sense | TCTGGATCGGCTACTGCAAC |
| | | antisense | AAGGGAAGCCCAGACATTGG |
| *hAdra2c* | NM_000683.4 | sense | GCCCGCTCTTCAAGTTCTTC |
| | | antisense | CCTTGCTTGCCCATTTAGGG |
