## Supplementary figures and images for "Sympathetic Nervous System Overactivation Induces Colonic Eosinophil-Associated Microinflammation and Contributes to the Pathogenesis of Irritable Bowel Syndrome"

### supplemental figure

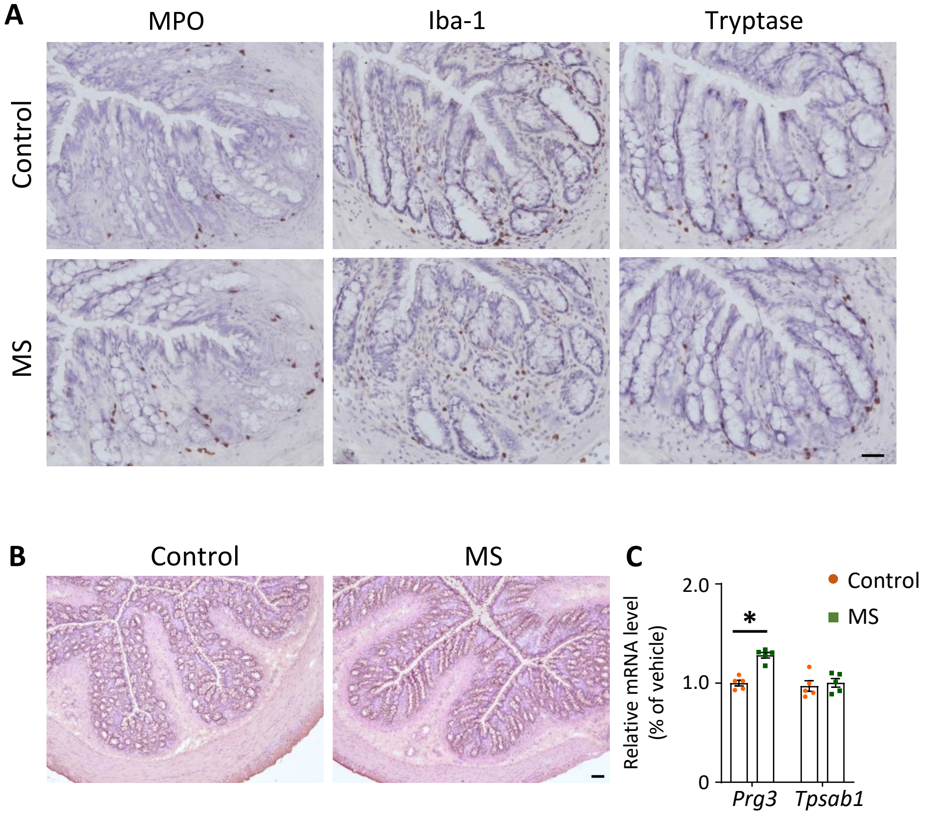

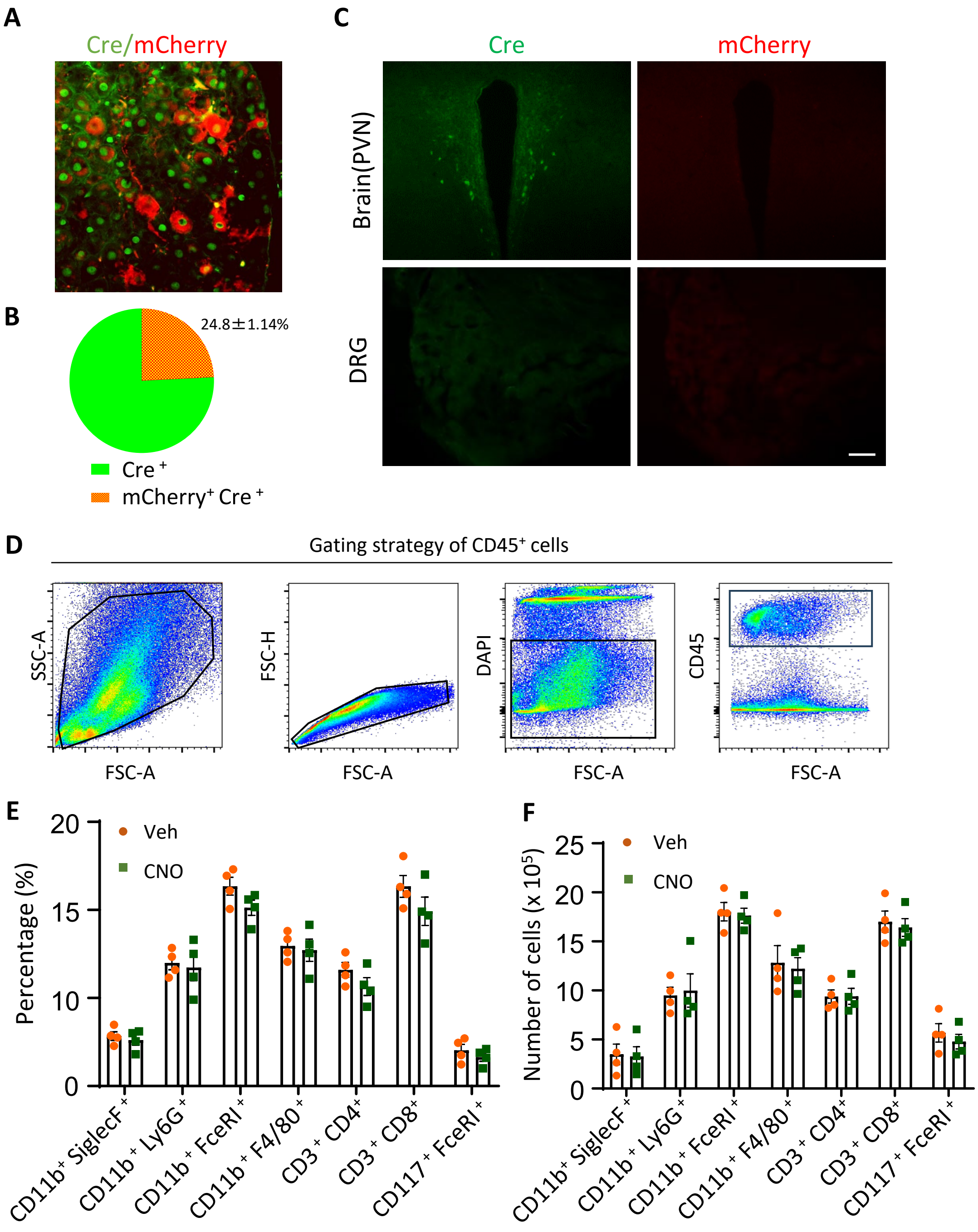

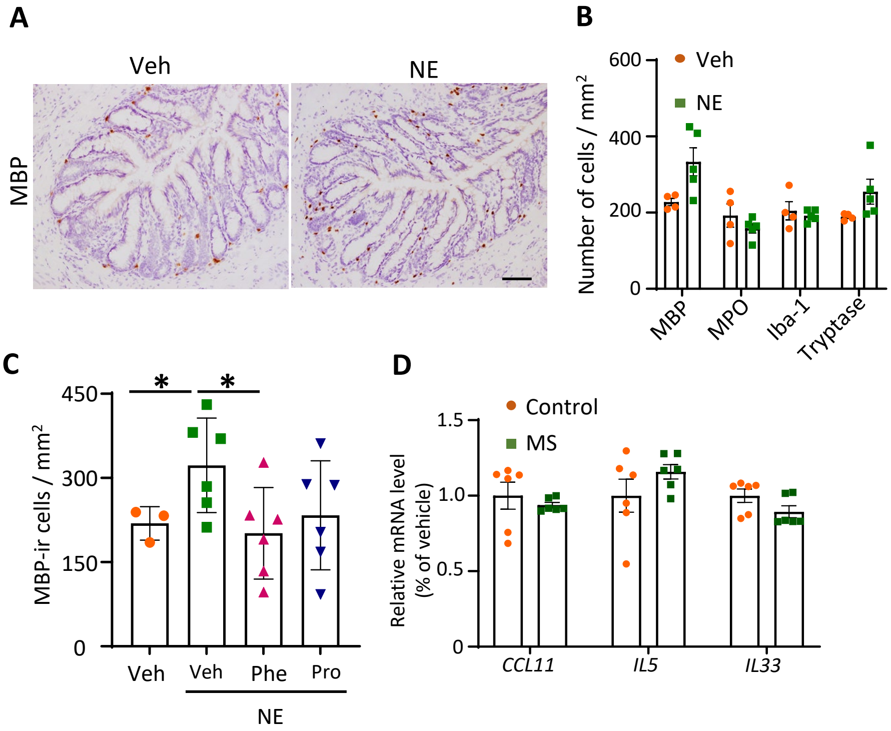

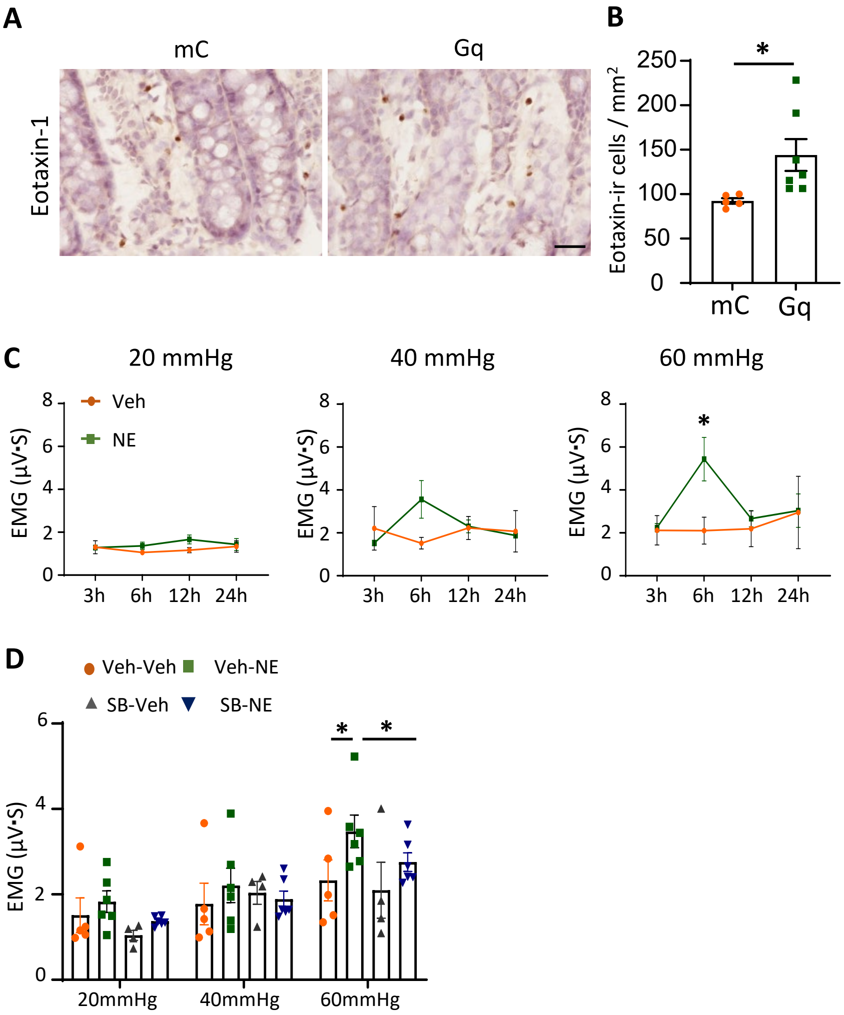
